# Raman Signal Denoising using Fully Convolutional Encoder Decoder Network

**DOI:** 10.1101/2022.01.07.475347

**Authors:** Irem Loc, Ibrahim Kecoglu, Mehmet Burcin Unlu, Ugur Parlatan

**Affiliations:** Bogazici University Physics Department, Istanbul, Turkey; Global Station for Quantum Medical Science and Engineering, Global Institution for Collaborative Research and Education (GI-CoRE), Hokkaido University, Sapporo, Japan

## Abstract

Raman spectroscopy is a vibrational method that gives molecular information rapidly and non-invasively. Despite its advantages, the weak intensity of Raman spectroscopy leads to low-quality signals, particularly with tissue samples. The requirement of high exposure times makes Raman a time-consuming process and diminishes its non-invasive property while studying living tissues. Novel denoising techniques using convolutional neural networks (CNN) have achieved remarkable results in image processing. Here, we propose a similar approach for noise reduction for the Raman spectra acquired with 10x lower exposure times. In this work, we developed fully convolutional encoder-decoder architecture (FCED) and trained them with noisy Raman signals. The results demonstrate that our model is superior (p-value < 0.0001) to the conventional denoising techniques such as the Savitzky-Golay filter and wavelet denoising. Improvement in the signal-to-noise ratio values ranges from 20% to 80%, depending on the initial signal-to-noise ratio. Thus, we proved that tissue analysis could be done in a shorter time without any need for instrumental enhancement.

## Introduction

Raman spectroscopy is a vibrational spectroscopic method invented by CV Raman in 1928.^1^ The method is commonly applied for molecular structure analysis in many fields and is very effective while working with biological materials since it is not affected by water absorption.^2^ On the other hand, Raman spectra are very weak in intensity compared to other spectral techniques. When the optical scattering losses are added to the weak intensity factor, the signal-to-noise (SNR) values obtained in the biological sample analyses become very poor unless high exposure time acquisition or complex data processing algorithms are used.^3–5^ As a particular case, Raman spectral mapping from tissue samples becomes highly difficult and time-consuming, since spontaneous Raman scattering cannot provide a meaningful signal. Resonance methods such as surface-enhanced Raman scattering (SERS) are helpful to enhance the spectra. However, in the case of SERS, a gold substrate, which interferes with Raman signals, is needed to enhance the signal. ^2^ Therefore, computational methods are highly favored in situations where a good quality signal is hard to obtain.

The common computational methods for Raman spectra denoising are the Savitzky-Golay (S-G) filtering^6^ and wavelet denoising.^7^ The S-G filter fits a polynomial function to sliding windows on the signal. The parameters of the S-G filter, i.e. polynomial order and the window size, must be optimized, and each signal may require a different choice of parameters. On the other hand, wavelet denoising benefits wavelet transforms to decompose the noisy signal. This method requires choosing optimal wavelets, thresholding techniques, and leveling.

Deep learning (DL) stands as a novel and promising approach in signal denoising.^8,9^ The recent studies illustrate that artificial neural network (ANN) architectures outperform the state-of-the-art techniques for Raman signal denoising, such as the S-G filter^10,11^ and wavelet denoising,^10,12^ in terms of noise reduction while preserving the peaks in the Raman signals. In these studies, Gaussian noise is added to the actual Raman signals with high exposure times ^10^ or computer-generated signals,^11^ then fed to neural networks for noise reduction. Based on our research, no study worked with real noisy Raman signals.

We propose a fully convolutional encoder-decoder network (FCED) for denoising the Raman spectra. Similar to deep image denoising algorithms, our method benefits from convolutional layers to extract the features of the input signal. The FCED architecture takes the noisy Raman signal acquired in a short period as the input. It learns to map the noisy signal to the output signal acquired in a longer period. Therefore, we aim to reduce the time required to scan the materials with Raman spectroscopy. Our ultimate goal is to analyze tissue samples using Raman spectroscopy in a feasible time range.

## Experimental Section

### Raman Experimental Setup

We built a custom setup for spontaneous Raman spectroscopy. The excitation source was a diode laser with a wavelength of 785 nm and output power of 300 mW. We focused this beam into the quartz cuvette using a lens whose focal length is 50 mm. For the powder measurments, we used the micro-Raman scheme, where the excitation beam is focused using a 10x (0.25 NA) microscope objective. We collected the Raman beam into the entrance slit of the portable spectrometer (Ocean QEPro Raman) using a multimode fiber.

### Data Acquisition and Preparation

We acquired the Raman spectra of the chemicals listed above in two different schemes: Micro-Raman (for powder and tissue samples) and normal spontaneous Raman (for liquid samples). The liquid samples were measured in a quartz cuvette (Hellma) which minimizes the fluorescence background from the sample. For the chemicals with high Raman activity like toluene, acetone, methanol, isopropanol, we adjusted the exposure time between 50-100 ms and the laser output power to about 50 mW for noisy measurements. We only increased the exposure time (between 500-1000 ms) for high-exposure measurements with these chemicals. On the other hand, we used full laser power (300 mW output) and 100 ms exposure time for the remaining samples while acquiring the low-exposure dataset. We set the exposure between one and three seconds for the high-exposure measurements with the remaining chemicals and tissues. We collected 1000 spectra for both low and high exposure measurements from each sample. The spectra had an autofluorescence profile that could affect the model performance. We removed this background by subtracting a polynomial curve from each spectrum. Figure 1a demonstrates the combined spectra of all chemicals in our dataset.

**Figure 1:**
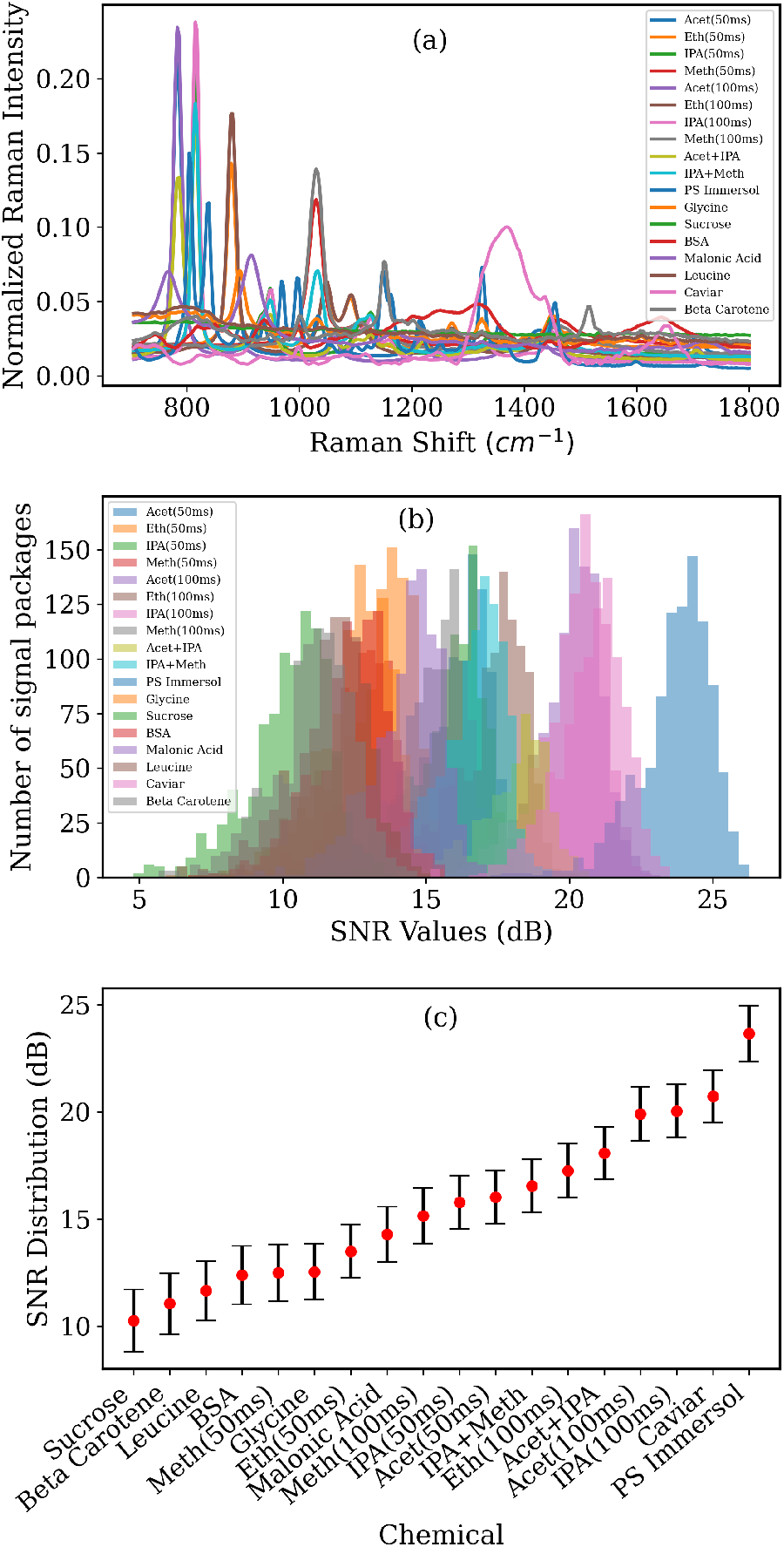
a)Combines spectra of all chemicals b) SNR distributions of each chemical c)Mean and standard deviation of SNR values

To train and test our architecture, we designed two experiments. For the proof of concept, we used Raman signals that are collected with high exposure times (between 500 to 3000 ms). Then, we artificially added previously measured white noise to these low noise signals. After adding the noise profiles to each chemical, we performed normalization using the *L*_2_ norms of high-exposure signals. Then, we subtracted the minimum value of the noisy signal from both high and noisy signal. With this procedure, we created a database of Raman signals with SNR values ranging from 10 to 25 dB. The SNR distribution of chemicals is shown in Figure 1b-c.

For the application part, we used Raman spectra of chemicals with both low and high exposure times. Each signal was divided by its acquisition time (in ms) for normalization. We developed a different approach than the previous part, where we applied *L*_2_ normalization to the data. Otherwise, we did not observe an overlap between high and low exposure signals.

We fed the low exposure signals into the FCED network as the input, whereas we passed the high exposure ones as the final output.

### Fully Convolutional Encoder Decoder Network Architecture

Deep learning is a general term, referring to the machine learning (ML) process using the multilayer (deep) ANNs. Convolutional neural networks (CNNs),^14^ which use convolution operations, are special types of ANNs. Convolutional layers are popular tools in many application areas, like computer vision and natural language processing. They use relatively fewer parameters than fully connected layers. They are good at capturing spatial features while being shift-invariant. These properties make CNNs powerful for signal processing.

On the other hand, the encoder-decoder is a well-studied architecture in ANNs. As the name suggests, the first half of the network encodes the input and reduces its dimensions. The middle layer that captures the encoded input is called the bottleneck. As the last step, the decoder part of the network tries to reconstruct the input signal from the encoded information. Encoder-decoder architecture can be built using fully connected layers or convolutional layers. Dimensionality reduction enables the network to capture the core information of the given input and eliminate the noisy parts. Since the noisy aspects of the input signal are removed in the bottleneck layer, encoder-decoder architecture stands as a strong candidate for denoising purposes.

In this study, we selected one dimensional fully convolutional encoder-decoder (FCED) architecture for denoising the Raman signals. We composed encoder blocks using two stacked convolutional layers with stride one and rectified linear unit (ReLU) activation and a maxpooling layer. We selected the initial filter size as 16 and doubled it at each consecutive encoder block. We formed the decoder blocks using two transpose convolution layers with stride one and ReLU activation, followed by an upsampling layer for signal reconstruction. Our network had three encoder layers and three symmetric decoder layers, and a final convolutional layer.

The schema for the encoder and decoder blocks and the entire architecture is shown in Figure 2. For each convolutional and transpose convolutional layer, kernel size was selected as three. The network was optimized using Adam optimizer^15^ of Tensorflow Keras backend.^16 1^

**Figure 2:**
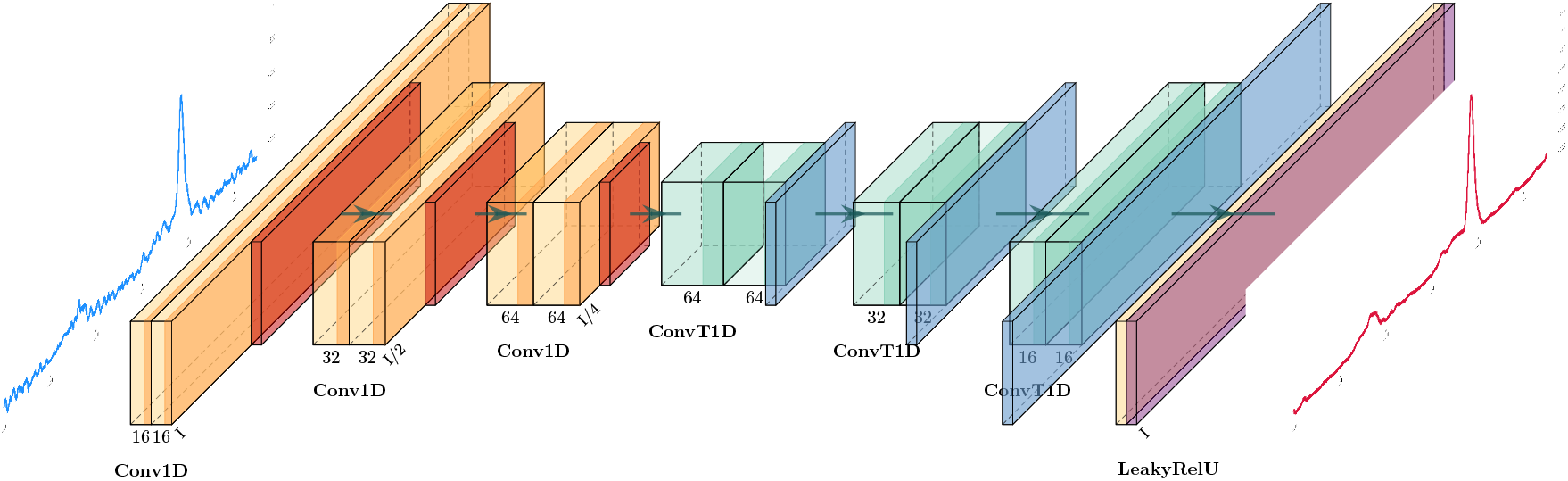
The architecture of FCED Network. Yellow and orange blocks represents the 1D convolutional layers (Conv1D) with ReLU activation followed by max pooling layer. Green and blue blocks are 1D transpose convolutional layers (ConvT1D) with ReLU activation followed by an up-sampling layer. Violet layer at the end represent the final 1D convolutional layer with activation LeakyReLU^13^.

### The Custom Loss Function

We take the idea of the non-standard loss function from Barton et al.,^11^ which is designed for preserving the peak fidelity, and we created our loss function. In both versions, the custom loss function combines the regular mean square error (MSE) loss with local MSE loss. Though, the definition and the parameters for local MSE loss differ. The equation 1 is the regular MSE loss, while local MSE loss is given in the equation 2 and 3. *S_ref_* denotes the reference noise-free signal, whereas *S_noisy_* is the noisy/denoised signal.

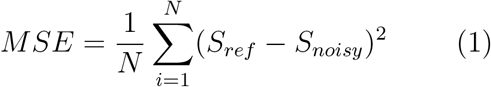

We selected the parameter *t* as the percentage threshold for the peaks in the signal. We discarded the points that are below the *t* percent of the maximum value. We only take into account the peak points, i.e. the points that passed the threshold, for the local MSE loss function.

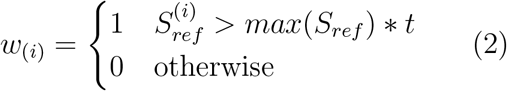

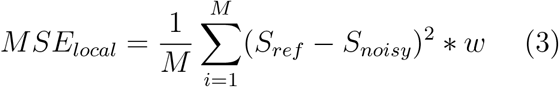

We did a series of experiments to determine the optimum percentage threshold t. Initially, we trained five different models with parameter *t* = 5%, 10%, 15%, 25%, 50%, respectively. Then, we tested their SNR improvement with four different chemicals. The results are presented in the Figure 3a.

**Figure 3:**
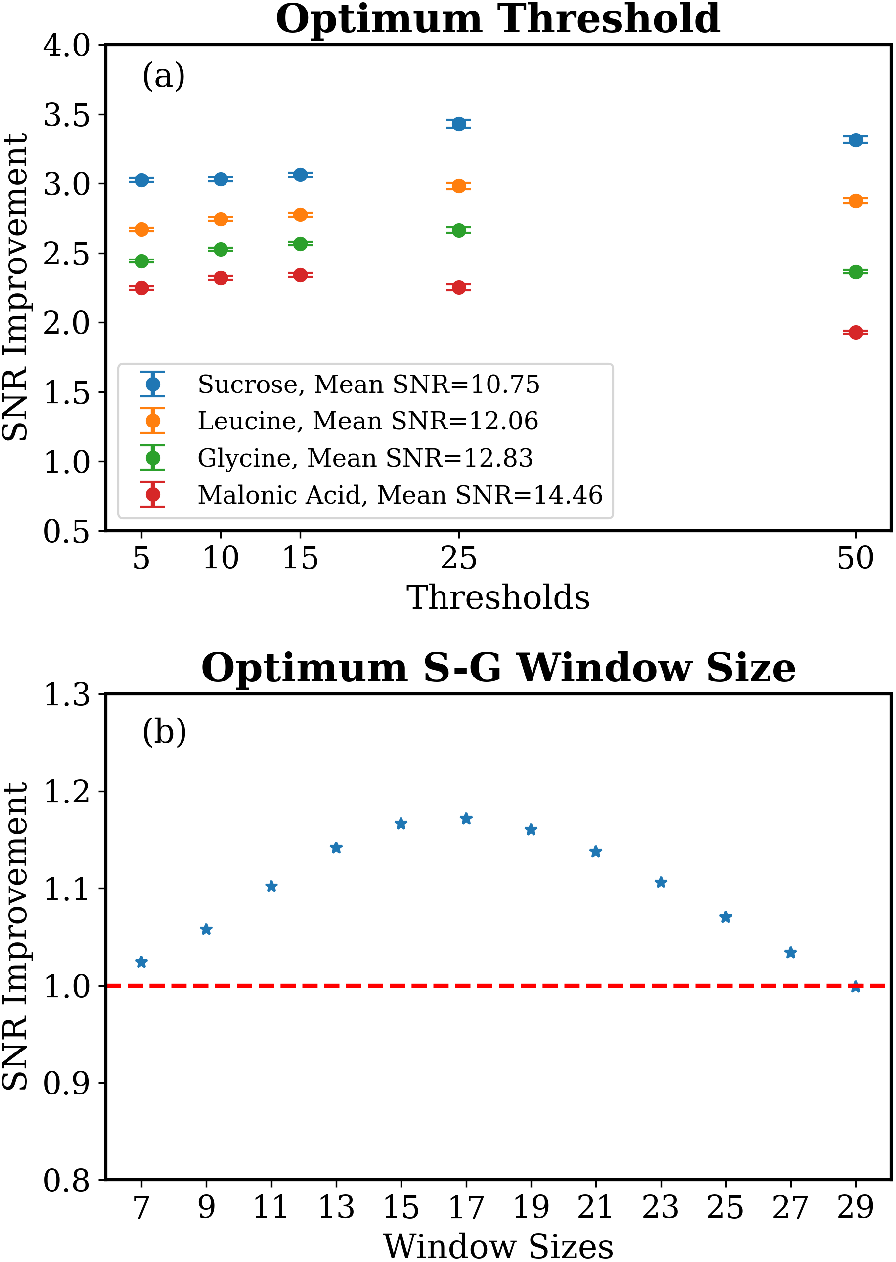
a)SNR improvements of different thresholds for the custom loss function b) SNR improvements of the S-G filter with various window sizes.

We observed that the increased threshold, to a certain level, had a positive effect on all chemicals. As seen in the Figure 3a, three out of four chemicals had the peak around 25% threshold value for their SNR improvement results. Therefore, we selected 25% as our final threshold value.

### Evaluation Metrics

Three evaluation metrics and their improvement after denoising are used to test the performance of our model. The first two are SNR and mean absolute percentage error (MAPE). For the third one, differences between the reference signal and noisy/denoised signal are weighted according to their intensity values and summed up. We named the metric as weighted absolute differences (WAD), and it is a measure of how well the peaks are conserved. The equations are provided below.

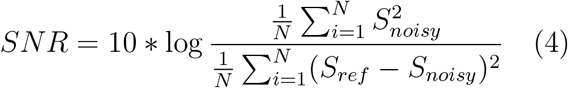

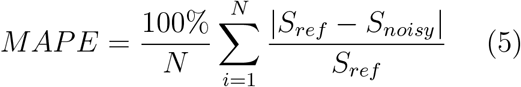

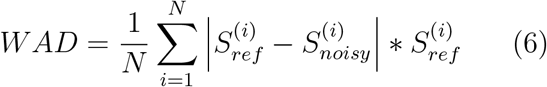

We tested the improvements by calculating the relevant metric value for both input (noisy) and output (denoised) signal, then calculating their ratio.

To compare our model with the classical denoising techniques, we need to determine the parameters of the S-G filters and the wavelet denoising. Based on our experience and observations, we selected polynomial order of the S-G filter as three. Then, we tried possible candidates for the value of the window size, and compared their (both global and local) SNR improvement scores. We decided that the optimum window size was 17, as demonstrated in the Figure 3. For the wavelet denoising, we used Daubechies 8 (db8) wavelet with level 3.

## Results and Discussion

### The Proof of Concept

The first part of our research was to study the noise reduction performance of our FCED model with the Raman signals that we artificially added noise. This part was the proof of concept of our work.

We trained our FCED model with both regular MSE loss and modified version of it, which we explained in the Methods section in detail. We compared the results of our model with the S-G filter and wavelet denoising techniques.

Figure 4 illustrates the results of our models and the other methods, on the Raman spectra of ethanol. Ethanol is selected as test compound and not included in our training set. Figure 4c-e are the box plots of SNR, MAPE and WAD distributions, respectively. Each box plot includes the noisy signal and denoised signals using the S-G filter, wavelet, FCED with MSE loss, and FCED with the custom loss. Figure 4a demonstrates an example denoising result with FCED with MSE loss, whereas Figure 4b is for the custom loss.

**Figure 4:**
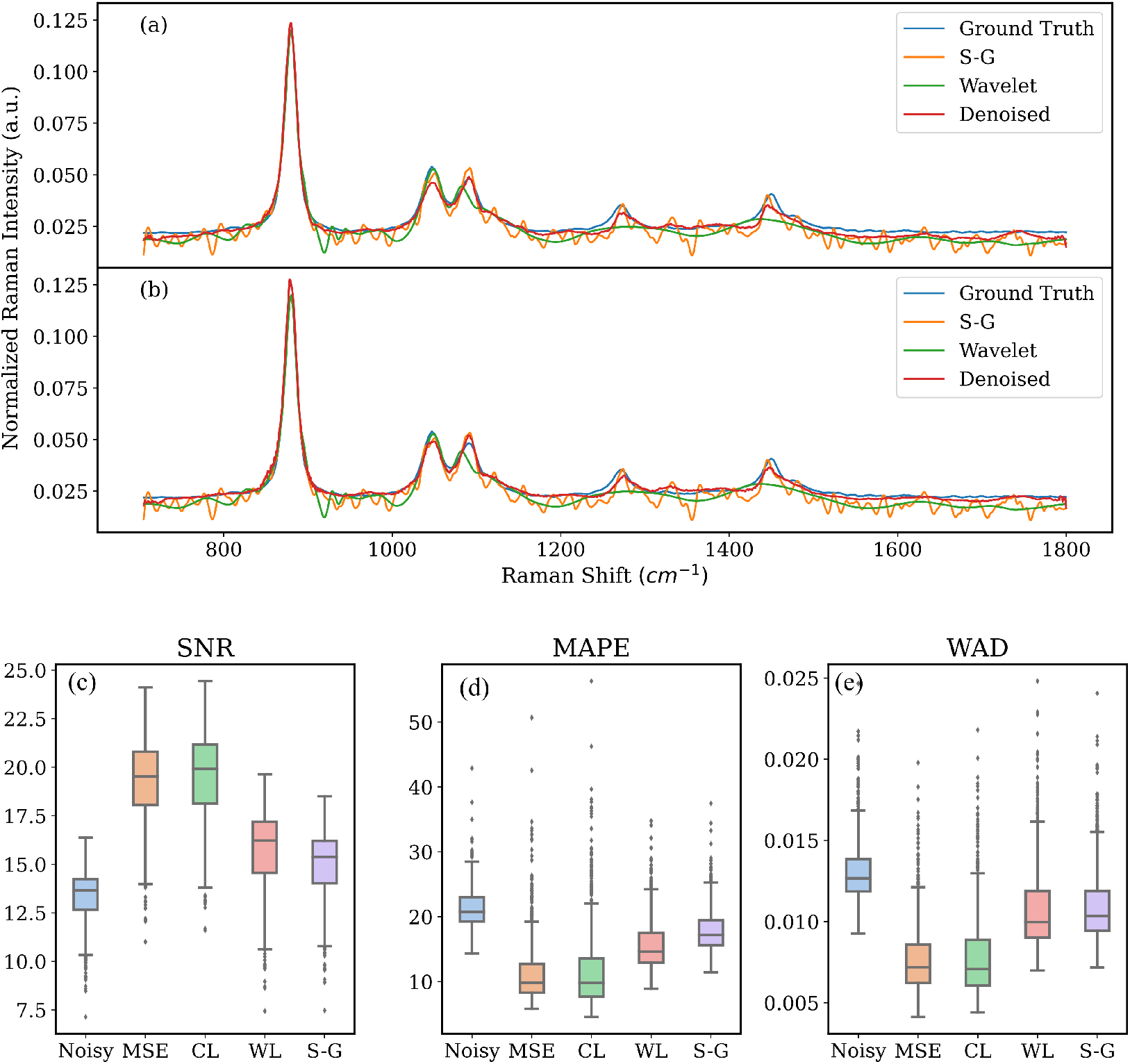
Denoising results of artificially noise added ethanol(500 ms) spectrum a)FCED model trained with MSE loss, b)FCED model trained with the custom loss, c)change in the signal-to-noise ratios, d)change in the mean absolute error (MAPE), d)change in the weighted absolute differences (WAD). Y-axes represent SNR (dB), MAPE and WAD values, respectively.

The performance of the FCED architecture trained with the custom loss exceeds other methods at peak fidelity (preserving the peaks). Nevertheless, we observed tiny distortions at the tails of the most prominent peaks, which we did not observe at the MSE loss (Figure 4a and b).

The Figure 5 demonstrates the mean and the standard error of the improvements of selected denoising techniques. We were inspired by Barton et al. and plotted our improvement graphs concerning the changing SNR values. We calculated the effect of the initial SNR value to the performance of our models. We found that increasing SNR lowers the performances of all methods. Regardless of the loss function, the FCED models are superior to the S-G filters and wavelet denoising for SNR values smaller than 20 dB. In addition, the custom loss is superior to MSE loss when initial SNR value is below 15 dB.

**Figure 5:**
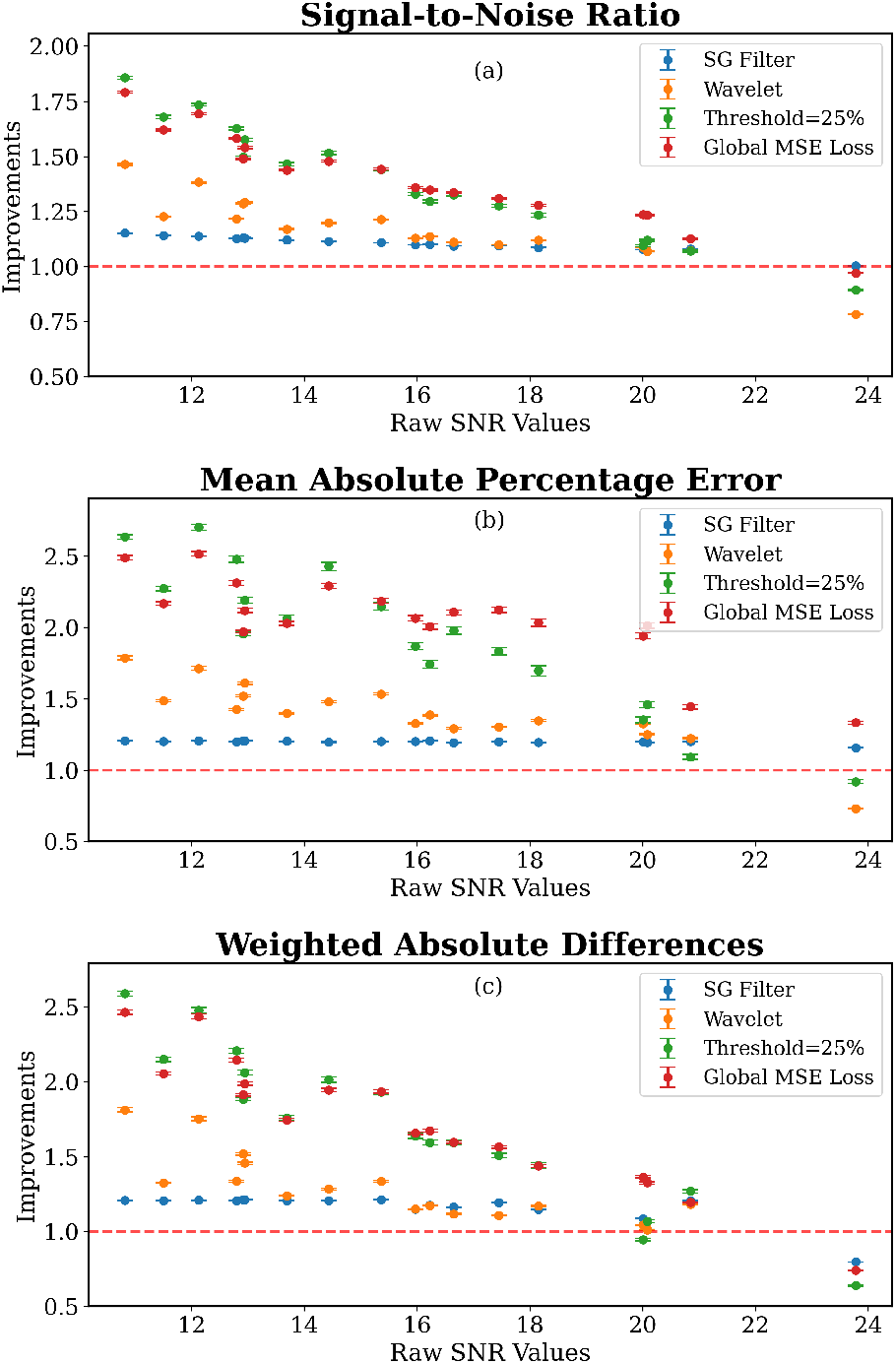
Mean improvements of different chemicals, which are represented in the x-axis by their initial SNR values. Figure demonstrates the results of the S-G filtering, wavelet, FCED model with global MSE loss and FCED model with the custom loss (with threshold 25%). a)SNR, b)MAPE and c) WAD improvements

This part of our study proved that FCED architecture, regardless of the loss function, outperforms the classical methods for Raman signal denoising at both peak fidelity and noise reduction.

### Practical Application

For the practical application of our approach, we trained our model using both low and high exposure Raman signals. Then, we tested our model with several compounds that are unseen by our model during training. Our test set included methanol, agarose, gelatin and atrium tissue sample. The mean (± standard error) of initial SNR values were 12.88(±1.35), 14.79(±0.95), 14.24(±1.03), 8.04(±1.13) for methanol, agarose, gelatin and atrium tissue, respectively.^2^

The spectra of test set and denoising performances of the S-G filter and FCED model was shown in the Figure 6. We did not include wavelet denoising results to the Figure 6 regarding its poor performance.

**Figure 6:**
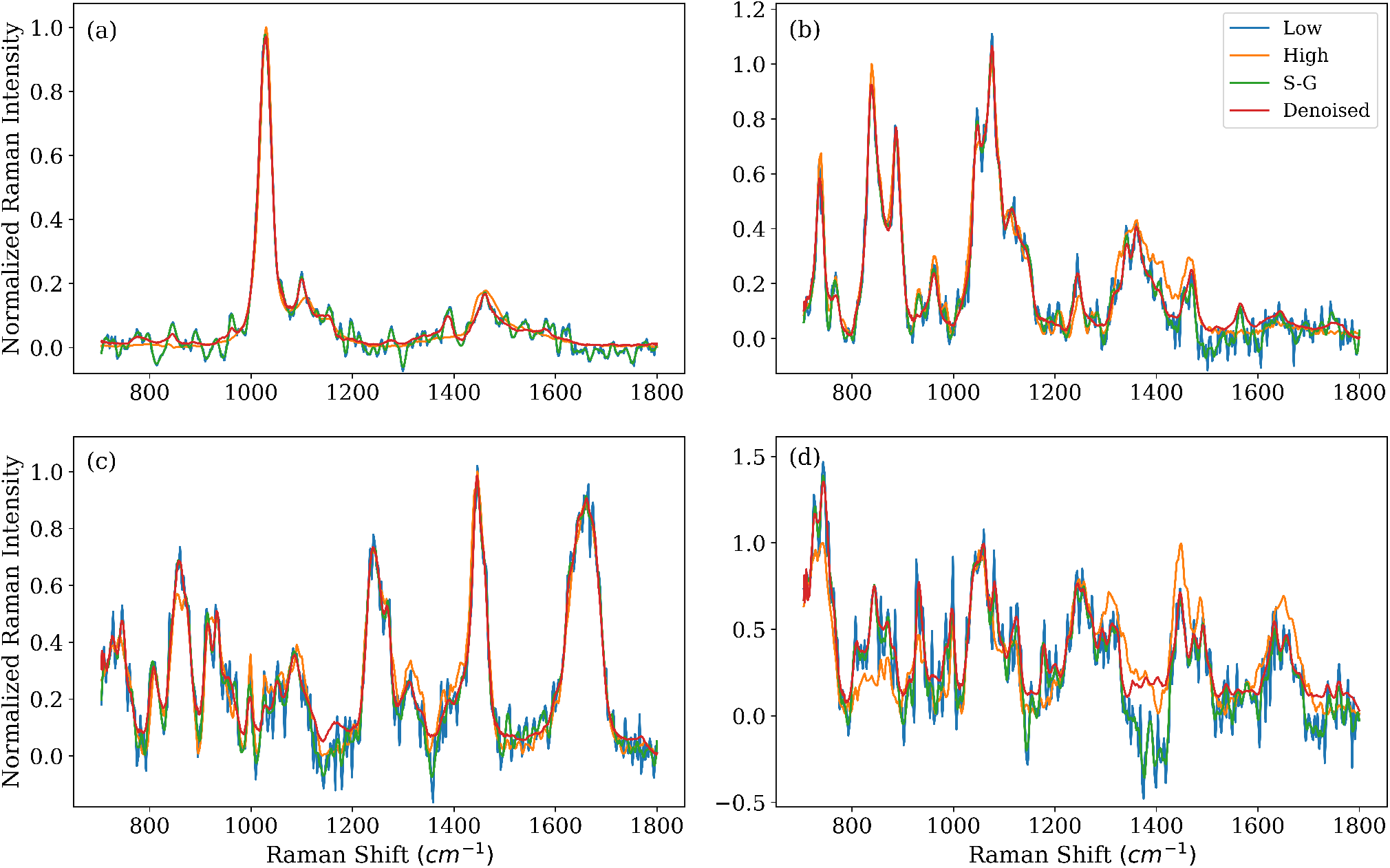
Ground truth(high), noisy(low), S-G filtered and FCED denoised spectra of a)methanol b)agarose c) gelatin and d) atrium tissue.

The mean SNR improvement score of the FCED model for the 997 atrium tissue signals is 1.342(±0.044), whereas the improvement of the S-G filter is 1.154(±0.001) and wavelet is 1.087(±0.001). When we conduct Student’s t test on the denoising performances of the S-G filter and FCED, we found that the p-value < 0.0001.

Our results indicate that using FCED architecture, we can recover a significant amount of the peaks in the high-resolution Raman spectra. The analysis shows that we could not recover some of the peaks with too low or too high SNR. However, one can overcome this issue utilizing a richer training set. On the other hand, since many times we only use the information from the significant peaks while doing peak analysis, this tool is still a candidate to be used in tissue analysis. Since we denoised the spectra that we acquired 10x faster, this denoising algorithm can speed up the Raman tissue analysis without a need for instrumental improvements.

## Conclusion

In this study, we developed a deep learning approach for Raman signal enhancement. We constructed the FCED model and trained it with different loss functions. We achieved peak preservation more accurately using the combination of global and local MSE loss as the custom loss. We were able to enhance the Raman signal that was acquired in a short time, better than the S-G filter and wavelet denoising techniques. In addition, we demonstrated that the FCED model could be used in any Raman signal whose SNR value is below 20 dB.

This study reduces Raman acquisition times by up to ten times, which in turn reduces tissue scanning time by the same amount. This improvement is an important step towards overcoming one of the biggest obstacles in point mappings, the inability to obtain a practical measurement time for medical applications.

## Acknowledgments

This study was supported by a grant from the Ministry of Development of Turkey (Project Number: 2009K120520).

